# HTSvis: A web app for exploratory data analysis and visualization of arrayed high-throughput screens

**DOI:** 10.1101/107821

**Authors:** Christian Scheeder, Florian Heigwer, Michael Boutors

## Abstract

The analysis and visualization of arrayed high-throughput screens (HTS), such as cell-based RNAi or small-molecule HTS experiments, requires specialized computational methods. Software packages such as the R/Bioconductor package cellHTS have been developed to support the analysis and are broadly used by the high-throughput screening community. However, exploratory data analysis and integration of screening results remains challenging due to the size of produced data tables in multi-channel experiments and the lack of user-friendly tools to integrate and visualize screening results. Here we present HTSvis, an R/Shiny open-source web application for interactive visualization and exploratory analysis of arrayed high-throughput data. Using a light-weight infrastructure suitable for desktop computers, HTSvis can be used to visualize raw data, perform quality control and interactively visualize screening results from single- to multi-channel measurements, such as image-based, screens. Input data can either be a result file obtained upon analysis with cellHTS or a generic table with raw or analyzed data from, e.g. a high-content microscopy screen. HTSvis can be downloaded from http://github.com/boutroslab/HTSvis.

## Introduction

Arrayed high-throughput screens (HTS) in high-density multi-well plates are a powerful method for target and small molecule discovery [1,2]. Automated technologies allow to screen tens of thousands of genetic or chemical perturbations, resulting in very large data sets. High-throughput screens can range in complexity from single-channel cell viability measurements [3,4], to multi-channel FACS [5] and multi-parametric imaging screens [6–8]. To analyze the resulting data sets, a range of statistical methods have been developed for processing, normalization and quality control of HTS data. Data then has to be further processed to identify significant perturbation and annotate identified ‘hits’ [9–13]. Open-source software for integrated statistical analysis using statistical languages such as R, have been developed [14–17]. While commercial software packages, e.g. TIBCO Spotfire, exist for visualization and exploratory data analysis on desktop computers, few open-source options are available, in particular for multiparametric imaging screens [18,19]. Thus, there is a need for software applications that are easy to install and straight-forward to use to explore and visualize large HTS data sets.

Here we present HTSvis, a scalable web-based application for the visualization of data from high-throughput screening experiments. Data can be uploaded in commonly used formats to store raw- and analyzed data, such as delimited files or RData stores. In particular, data analyzed with the widely used cellHTS package be directly loaded for interactive visualization. In addition, the option to upload data in a generic tabular format, allows flexibility to visualize data of other assay types (e.g. multiparametric high-content data). HTSvis is designed as an application with a user-friendly interface using a *shiny* implementation of R, which runs in all standard web-browsers. User-interfaces facilitate data input and data views making HTSvis applicable for a broad range of uses. The interactive data presentations provide a compact, yet comprehensive view on the data as the entire data set can be browsed on interactive surfaces for plots and tables.

## Installation and data input

HTSvis is implemented as a web-application based on the R framework *shiny* (http://shiny.rstudio.com/) and is distributed as an open-source R package [20]. The HTSvis package is available through a GitHub repository (https://github.com/boutroslab/HTSvis) and can be installed directly from the R or RStudio console. Instructions for loading the package can be found on the github repository. After loading the package, the application is launched by calling the HTSvis function as follows *HTSvis()*. Once the function is called, the application will automatically open a window in the user’s default web browser. HTSvis is accessible to the users with minimal knowledge of the R language. Data upload and further usage are controlled via the user-interface and do not require R programming skills. HTSvis was developed with the aim to provide a performant web application, which can be used on local computers rather than a web-service where data should be submitted to an external server. The R based shiny framework provides a robust way to share web applications for local usage. The platform consistency between the R/Bioconductor cellHTS package [14] and HTSvis further helps streamlining data analysis and visualization (Figure 1). The application is organized in a tab-panel structure with five tabs (Figure 2A) including the *“Data Input”* tab and four tabs with plots and tables. When the application is started, the *“Data Input” tab* is shown and all other tabs are initially inactive. Data input is managed via a file importer, which allows choosing the input data from a local source. The input data should be in a tabular format as detailed in Supplementary File 1. For HTS data analyzed with cellHTS, the summary table *(“topTable.txt”)* of the analysis provides all required information and can be uploaded without further modification. Single and dual channel experiments analyzed with cellHTS can be explored and visualized with HTSvis (see also the documentation of the cellHTS package for further information concerning single and dual channel measurements). After data input, columns containing the required annotation such as the *well* and *plate* allocation are chosen from drop down lists containing all column names. Alternatively, the user has the option to upload data in a generic tabular format with certain requirements concerning the structure of the data table. Importantly, the number of measured channels per well is not limited which allows to load multiparametric data sets as from automated microscopy or flow-cytometry experiments. Detailed information, including a user manual, can be found on the application’s help page which can be excessed after launching the application.

**Figure 1:**
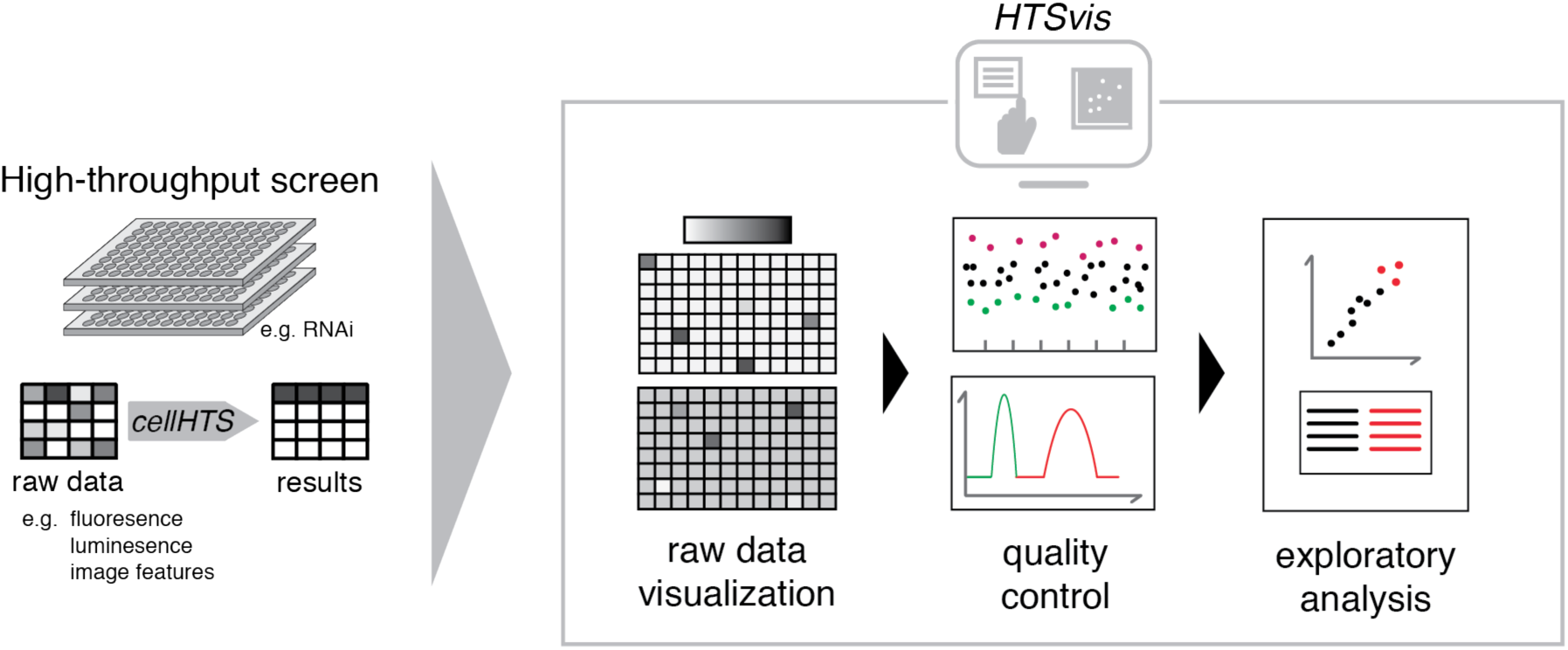
Workflow diagram for visualization and exploratory analysis of arrayed high-throughput screens. Raw data or statistically analyzed data from various arrayed screening formats, ranging from single channel readouts to image features in 12- to 384-well plates, in tabular formats can be loaded into the application to facilitate the identification of experimental artifacts, quality control checks and the identification of hits by interactive data representations.

**Figure 2:**
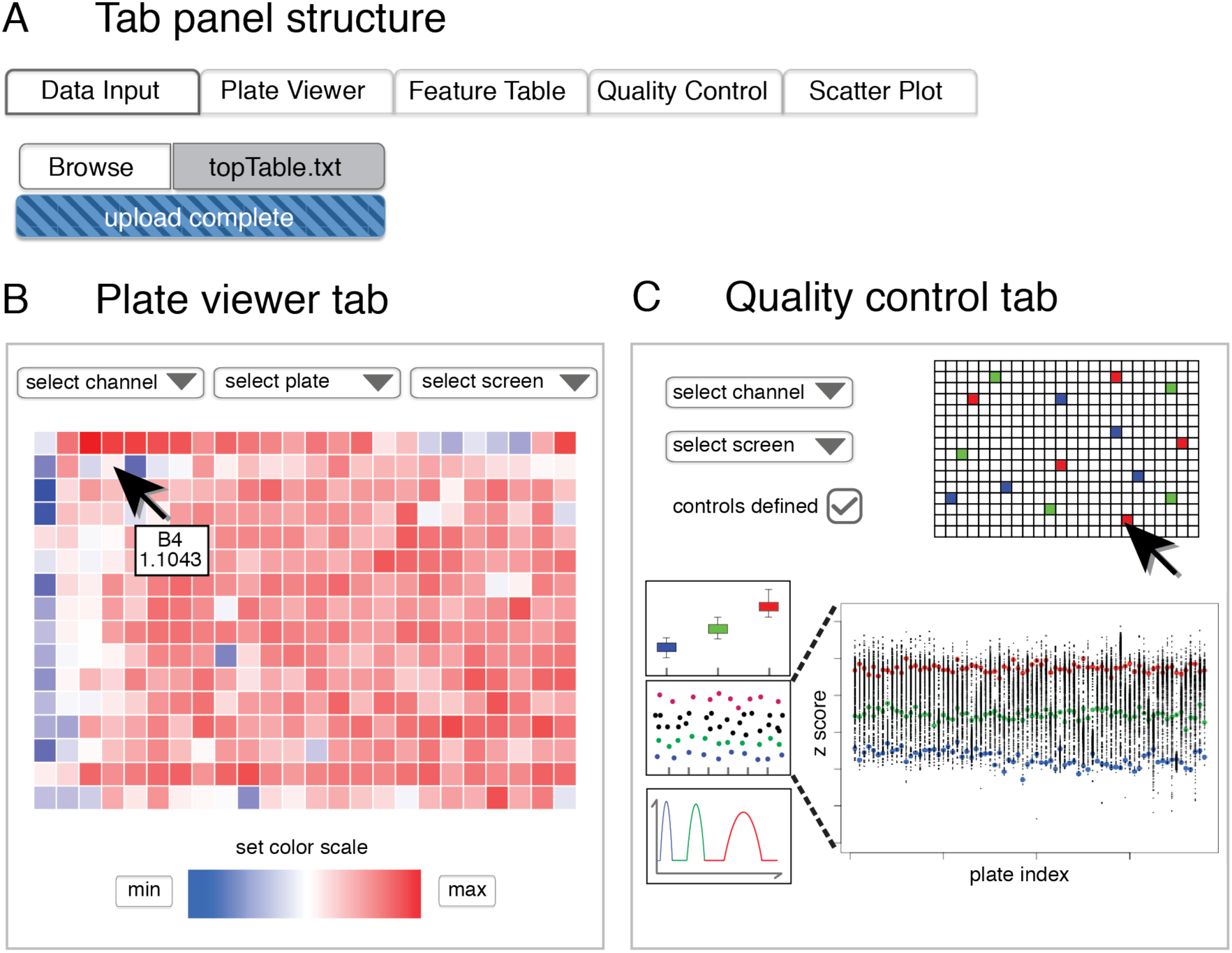
Schematic illustration of the user interface for interactive raw data and quality control visualizations. A) The tab panel structure and an automated file input provide a user interface for quick and intuitive data browsing. B) Multi-well heatmaps with a custom color scale and a tooltip help to identify experimental artifacts. The plate and feature to be shown for each heatmap are selected from drop down lists. C) The quality control tab provides informative plots per experiment based on user defined control wells.

## Interactive data exploration

After data input and selection of the columns with the *well, plate and experiment* annotation, four tab panels with data representations allow the user to interactively explore and asses the data. Due to the tab panel structure, a simple mouse click is required to switch between the tabs and a download handler is included for all plots and tables.

## Spatial plate analysis: Plate Viewer tab

A panel with four heatmaps is placed in the plate viewer tab. Each heatmap represents the data for one multi-well plate arranged and visualized in the assay format (e.g. 384-well plate format). For each of the four heatmaps, the plate and channel to be plotted are selected from drop down lists (Figure 2B). This allows the user to browse the data set and visually identify experimental artifacts, such as edge effects, and to compare specific plates within the data set, for example biological replicates. We designed a panel of four heatmaps to further compare performance of plates between channels or conditions, e.g. two biological replicates for each of two channels. Due to the chosen size of the plots, four heatmaps can be displayed aside on most monitors. The minimum and maximum values covered by the color scale can be adjusted manually for each heatmap. Missing or flagged data points (represented by the NA symbol in the R language) are indicated by black color. A tooltip, which is activated by hovering over the heatmaps, has been implemented. This additional feature provides accessible information about the measured value and annotation (e.g. perturbation reagent) per well.

## Data browsing: Feature Table tab

Based on the tabular format resulting from the cellHTS analysis (or other data from arrayed HTS experiments in a tabular format), we included a representation of the data as a table. Filters for each column of the interactive table allow rapid and intuitive filtering and sorting. The so filtered table can be exported as a *.csv* file. Each row of the data table contains the measured values of one unique well from one plate within the data set. If more than one measured value is assigned to each well (e.g. a dual channel luciferase assay), a hierarchically clustered heatmap can be created. Data from wells that should be included in the heatmap are dynamically selected and unselected by clicking on the corresponding rows of the data table. This feature is especially powerful for high-content data with per-well readouts presented by multi-dimensional feature vectors.

## Assessing screening quality: Quality Control tab

This tab is designed to facilitate quality assessment for experiments based on control samples. Data quality and integrity are assessed based on defined control populations for which a phenotype effect is expected. This strategy is broadly used by the screening community [10,11]. The behavior of up to three control populations (positive, negative and non-targeting controls) can be investigated. The controls are defined by their well position and are selected/unselected by clicking on a multi-well plate layout. Based on the user-defined control populations, three plots including a scatter plot, a box plot and a density plot (based on Kernel-density estimation) are provided to evaluate data quality and integrity. The box plot and density plot are useful to summarize how clear the control populations are separated over entire experiments. In addition, a numerical quality metric for the statistical effect size (the Z’-factor) is given in the legend of the density plot [12]. Both plots therefore provide data visualizations to estimate the statistical effect size and performance of the assay type. The scatter plot complements the summary statistics provide by the box- and density plot by adding information about measured values of individual plates. Identified outlier plates can then for example be investigated in detailed manner by switching back to the plate viewer tab. All quality control plots are immediately refreshed as the control populations are changed. The Measured channel and experiment for which the quality control plots should be created are chosen from drop down lists (Figure 3A).

**Figure 3:**
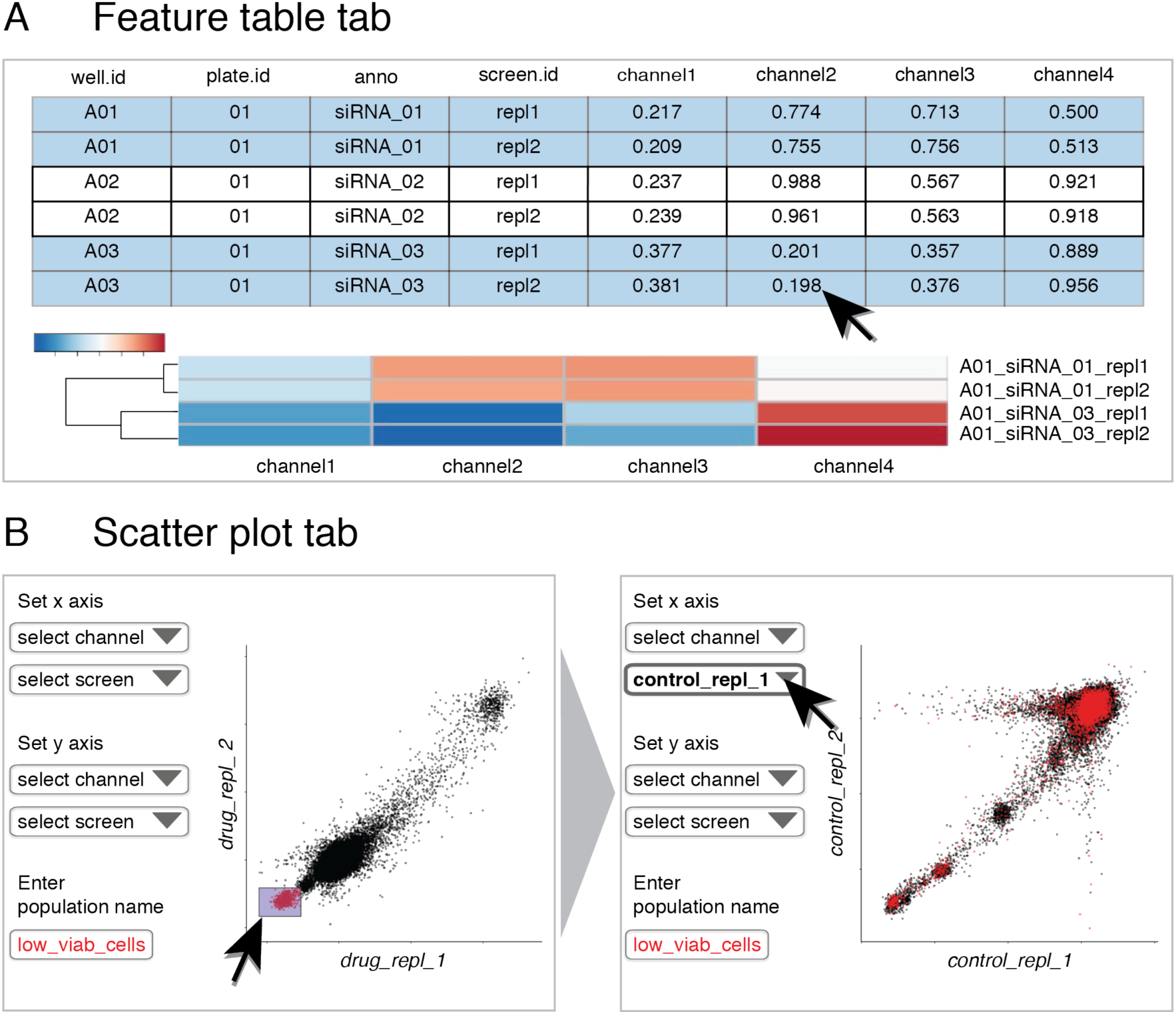
Interactive data representations facilitate exploratory data analysis. A) The feature table tab provides a table that can be searched and filtered for download. A reactive heatmap can be created to which rows are dynamically added and removed by selecting and de-selecting them in the table above. B) The scatter plot tab allows to assess the correlation between experiments and facilitates the identification of hits. Subsets of measurements can be grouped and colored with a mouse drag-and-drop. Brushed data points can then be compared between experimental conditions.

## Data interpretation: Scatter Plot tab

The scatter plot tab is a visualization tool for quality control and interpretation of the data. The central element is a scatter plot with two experiments plotted against each other as data series in a Cartesian grid. Such plots are helpful to evaluate the correlation between replicates and thereof judge the reproducibility of an experiment. Which experiment and measured channel the data should be plotted for is defined by drop down menu selection. We implemented an additional feature to brush data points by simply drawing a rectangle on the plot area. Data points inside the drawn rectangle can now be assigned as a subpopulation with a population name defined by the user. Multiple populations can be created in this way; each will be marked with a different color. Once a population is defined, all data points assigned to this population will be marked. Furthermore, the identity of those data points can be accessed via a table containing the well and plate ID of each marked point. This approach allows testing hypothesis and identifying measurements of interest associated with such, for example perturbations that induce a loss-of-function phenotype. Importantly, the labeling of data points is linked to the well and plate position and is preserved when the data input is changed. Assuming a fixed plate layout, differential effects between conditions (e.g. control and drug treatment) can be identified using the scatter plot in combination with the population manager (Figure 3B).

## HTSvis for data in a generic tabular format

All functions of the app described above are also available when the data input is a generic table and not a result file from an analysis with cellHTS. Given that the table has the correct format as described in Supplementary File 1, input data can be from any kind of arrayed screening experiment. The application supports 12-, 48-, 96- and 384-well plate formats. This flexibility opens the possibility to visualize raw data as well as data analyzed with an optional method.

## Conclusions

HTSvis is a web-application to explore and visualize data from arrayed high-throughput screens. Data from screening experiments that have been analyzed with cellHTS can be directly imported. The application is easy to install as an R package and launched on local computers from the R console. Various visualizations that are helpful for the assessment and interpretation of data from arrayed high-throughput screens are included. The interactivity of plots and tables provides a great advantage compared to the handling of individual files and programming scripts, for example one for each plate. Ease-of-use is the main characteristic of the graphical user-interface to facilitate user-friendliness from data input to data visualization. The interactive data representations further provide a versatile tool for exploratory data analysis filling a yet unmet need in the high-throughput screening community.

## Methods

HTSvis has been developed as a web application using the R shiny framework with R version 3.3.2 and shiny package version 1.0.0. The graphical user-interface was designed as a multi tab page with a fixed width of 1048 pixels. All calculations and data arrangements are implemented using the R language. Plate heatmaps in the pate viewer tab are implemented with the ggvis package (http://ggvis.rstudio.com/). For each heatmap a color scale is spanned individually between the minimum (blue) and maximum (red) value with 500 color steps in between. The interactive table in the plate viewer tab is implemented using the DT package, an R interface to the *DataTable* Javascript library (datatables.net). For the reactive heatmap of clicked rows in the feature table tab the *heatmap.2* function from the gplots package with the default complete linkage cluster algorithm was chosen. The density distributions for plotting in the quality control tab are computed as Gaussian kernel density estimates. The boxplot is created using the R base graphics package with the data points of control values plotted separately.

## Acknowledgements

We thank Luisa Henkel for helpful suggestions and comments on the manuscript and the Boutros lab for discussions.

**Funding**

Work in the lab of M. Boutros is in part supported by an ERC Advanced Grant.

